# Do invasive tetrapods conserve their climatic niches?

**DOI:** 10.1101/2024.03.28.587128

**Authors:** Biswa Bhusana Mahapatra, G. Ravikanth, N.A. Aravind

## Abstract

Invasive species are the second biggest threat to biodiversity after habitat fragmentation. The species list was collated from the Global Invasive Species Database (GISD) and the Centre for Agricultural and Bioscience International (CABI), and the species occurrence data were downloaded from the Global Biodiversity Information Facility (GBIF). This study examines the niche dynamics of 152 invasive tetrapods and finds that 60% of the species have very low or low niche overlap across the native and introduced regions. There are species with very high niche overlap, such as the Cane toad, Brahminy blind snake, Eurasian collared dove, Whitehead marmoset and Asian house shrew. Around 30% (=46) of species are showing considerable niche expansion. Similarly, 21 species (5 amphibians, 6 reptiles, 6 birds and 4 mammals) show no expansion in the introduced region. The introduction pattern presents that 46 invasive tetrapods are native to Asia, whereas 43 of them are introduced to North America.

## Introduction

Biological invasion is one of the greatest threats to biodiversity, ecosystem services, human health, and livelihoods (Lockwood et al., 2013; Early et al., 2016; Brondizio et al., 2019). Despite regulatory efforts, the global introduction of species continues to rise steadily (Seebens et al., 2017), with predictions indicating sustained high rates in economically developed countries due to unfolding climate change, intensification of tourism, and increased trade in plants and pets (Early et al., 2016; Seebens et al., 2017; Ribeiro et al., 2019; Seebens et al., 2021).

The intentional or unintentional introduction of invasive vertebrates, facilitated by the expanding pet trade and global movements, exacerbates the challenges associated with invasive species. Concurrently, climate change significantly influences the distribution patterns of both native and introduced species. Climate change is a pivotal factor shaping the future movement of alien species (Peterson et al., 2008; Bellard et al., 2013). Some previously innocuous alien species become invasive due to climate change, enhancing their competitive advantages over native counterparts (Pyšek et al., 2020).

Invasive species, typically possessing broad ecological tolerances and adaptability to diverse conditions, exhibit rapid spread and colonisation of disturbed habitats. This adaptability enables them to thrive in extreme climate conditions, outcompeting native species (Diez et al., 2012). However, the impact of climate change on invasive species remains a subject of debate, with modelling studies yielding divergent conclusions— some suggesting an increase in the area invaded, while others indicate contraction in their distributions (Gallardo et al., 2017; Bellard et al., 2018).

The investigation into whether a given species experiences a shift in its climatic niche between its native and alien ranges has become more significant in recent times (Bates et al., 2020; Mahapatra et al., 2023; Yang et al., 2023). This is not only because of the implications for niche-based predictions regarding the spatial transferability of alien species but also because of concerns over global change (Guisan et al., 2014). The extant body of literature has generated contradictory findings regarding the proposition that alien species retain their native climatic niches in the alien range, invaded range, or home away from home (Aravind et al., 2022). While some studies provide support for this notion, others contest it, indicating the invasive species shift its range or find a new home (Broennimann et al., 2007; Petitpierre et al., 2012; Early & Sax, 2014; Liu et al., 2020; Aravind et al., 2022). The consequence of this is a contentious discussion regarding the significance of niche conservatism in the field of biological invasions.

A recent study by Liu et al. (2020) assessed more than 400 alien species and showed that during the invasion process, the majority of species tend to maintain their niches, with few changes occurring in the climatic niches within alien ranges. Notably, an expansion of niches was observed in certain taxa native to the island (Liu et al., 2020; Stroud, 2021). This observation implies that niche transitions may be more prevalent among island endemics throughout the invasion process.

Tetrapods, a diverse group of vertebrates, serve as an ideal study subject for understanding invasion patterns due to their extensive human-induced movements and widespread distribution. Unlike birds and mammals, herpetofauna, comprising amphibians and reptiles, exhibit smaller ranges, with temperature significantly influencing their distributions in these cold-blooded organisms. Investigating how these vertebrates respond to native and introduced niches provides valuable insights. This study aims to (a) estimate the introduction of invasive tetrapods across continents and (b) uncover niche dynamic patterns across 152 invasive vertebrates within the categories of amphibians, reptiles, birds, and mammals. Findings from this research will enhance our understanding of regions with high invasiveness and responses to climate change and support the development of actionable management plans for invasive species containment.

## Materials and Methods

### Species data

For the present study, the species were selected from the Global Invasive Species Database (www.issg.org) and the Centre for Agriculture and Bioscience International (CABI, www.cabi.org till December 2020). The distribution data for all species were downloaded from the Global Biodiversity Information Facility (GBIF, www.gbif.org) and CABI between July 2018 and June 2020. The species list was further reduced to 152 (15 amphibians, 29 reptiles, 44 birds, and 64 mammals) for niche dynamics analysis based on distribution data availability in both native and introduced ranges. For all species used in the present study, native and introduced ranges were collated from various sources, primarily from CABI and ISSG databases.

### Niche dynamics

We used nineteen bioclimatic variable layers from WorldClim (www.worldclim.org) for the current scenario (ver 2.1) with a spatial resolution of 2.5 minutes (Fick and Hijmans, 2017). Principal Component Analysis (PCA-env) estimated the niche overlap across the native and introduced ranges (Broennimann et al., 2012). The spatially thinned rarefied points were segregated into native and introduced regions. We used the ecospat package ver 3.2 in R studio ver 4.1.0 to assess the Schoener’s D index (niche overlap), Expansion (E), Unfilling (U) and Stability (S) (Broennimann et al., 2021; R Core Team 2018). We also performed niche equivalency and similarity tests to compare the native and introduced regions. The niche overlap index varies from 0 to 1, with zero being no overlap and one being complete overlap (Rödder and Engler, 2011).

## Results

Our result shows that a maximum number of 46 species have been introduced from Asia, followed by North America with 40 species. Ninety-seven species were introduced to multiple continents, and about 43 species have been introduced to North America (Fig. 1).

**Figure 1:**
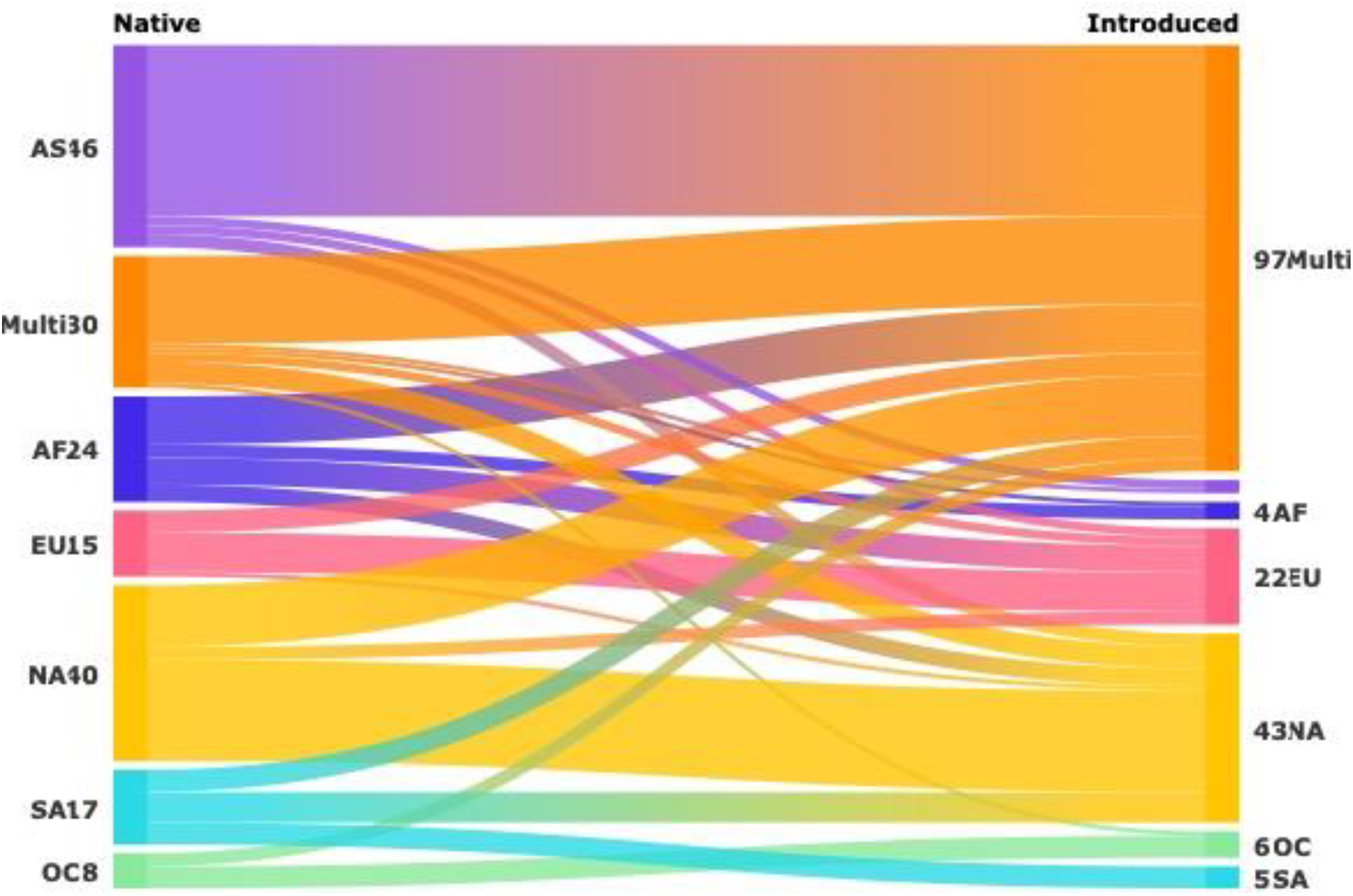
Sankey diagram shows the introduction of tetrapod species across different continents

### Niche dynamics

The overall trend of niche overlap indicates that more than 60% (92 out of 152 species) showed very low (49) and low (43) niche overlap (Fig. 3). Only five species, i.e., *Rhinella marina* (amphibian), *Ramphotyphlops braminus* (reptile), *Streptopelia decaocto* (bird), *Callithrix geoffroyi*, and *Suncus murinus* (mammals) showed very high (>0.80) niche overlap (Table 2). The equivalency test is highly significant, i.e., p-value <0.05 for 56% of mammals (36 out of 64), 75% of birds (33 out of 44), 40% of amphibians (6 out of 15) and 24% of reptiles (7 out of 29) (Table 2). This means the distinction between the native and introduced ranges is more in birds than in herpetofauna. The niche expansion varies from 0 to 0.86 in amphibians, 0 to 1 in reptiles, and 0 to 0.99 in birds (Fig. 2). We found zero niche expansion for five amphibians, six reptiles, six birds and four mammals (Table 2). The mean expansion (E) is lowest for birds and highest for mammals, i.e., 0.24 and 0.40, respectively (Fig. 2). Similarly, the lower mean unfilling (U) for birds is 0.36 and the highest for reptiles is 0.70 (Fig. 3C). The one-way ANOVA suggests there is a significant difference in both unfilling (F=6.77, p=0.000) and niche overlap (F=5.34, p=0.0016) among these taxa but no significant difference in niche expansion (F=2.24, p=0.086) (Table 3). 46 (∼30%) out of 152 invasive tetrapods show niche shift, i.e., (E > 0.5).

**Table 1:** ANOVA of niche expansion, niche unfilling and niche overlap among tetrapods.

|  | F | P-value | F crit |
| --- | --- | --- | --- |
| Expansion | 2.242 | 0.086 | 2.666 |
| Unfilling | 6.768 | **0.000** | 2.666 |
| Niche overlap | 5.344 | **0.002** | 2.666 |

**Table 2:**
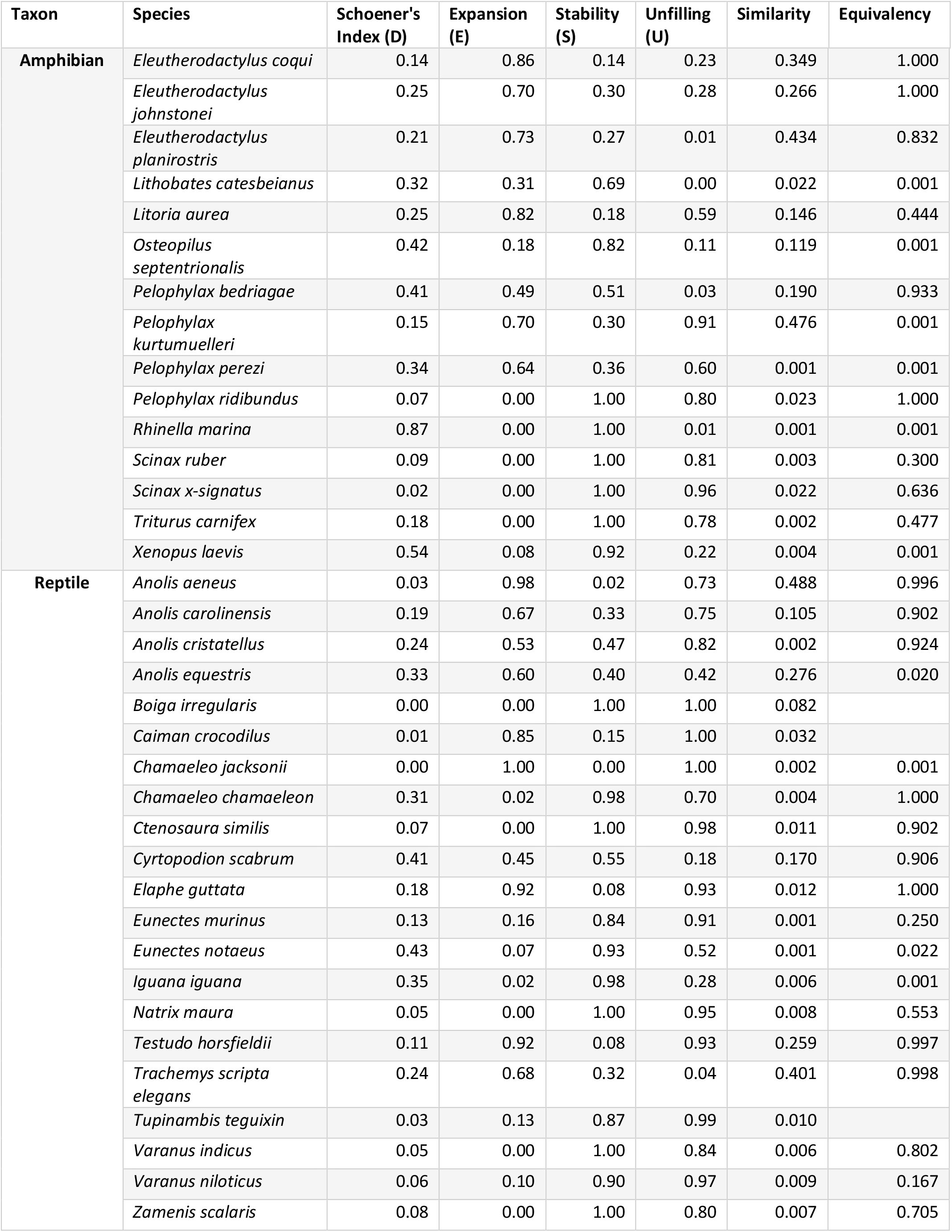
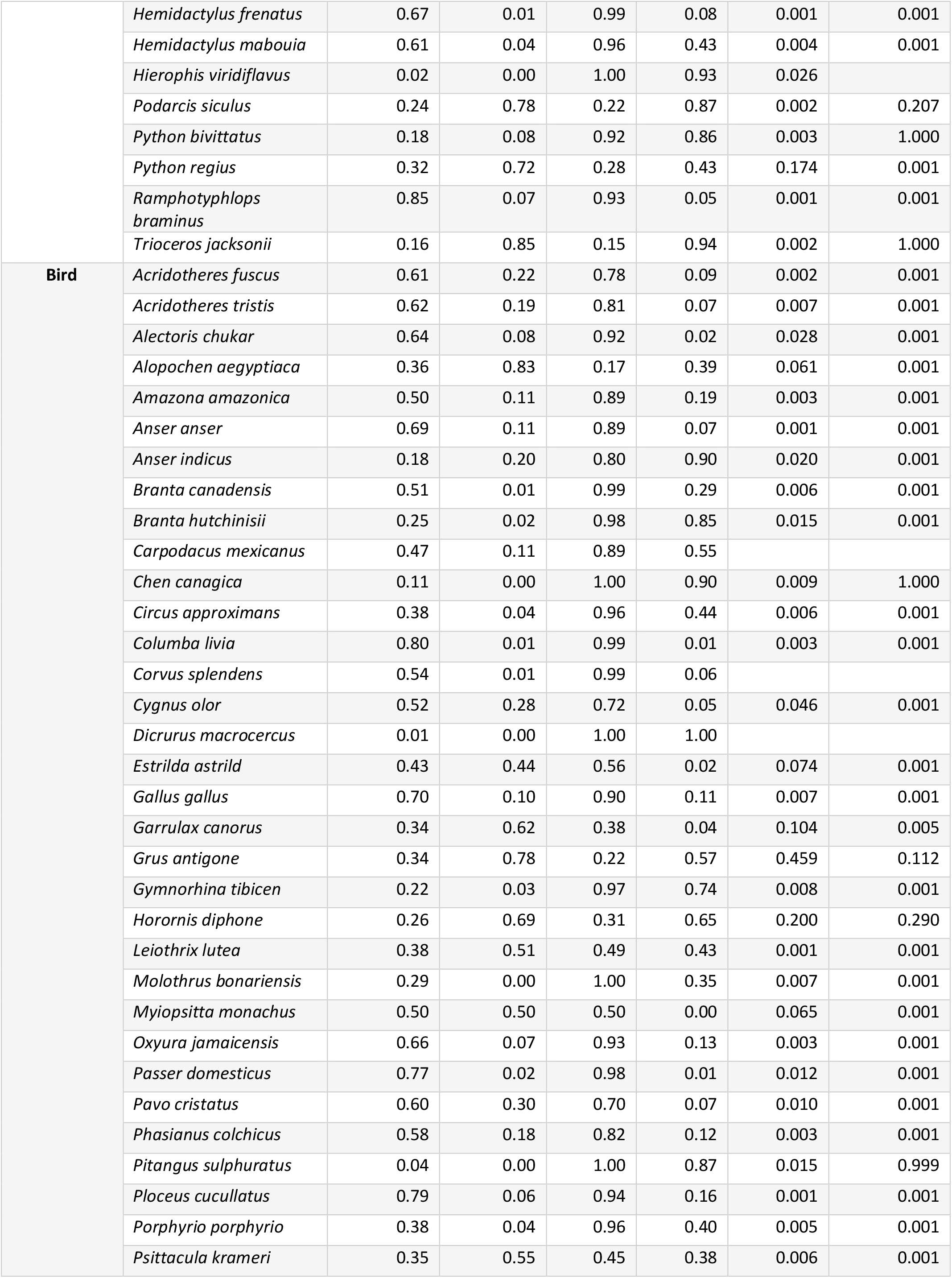
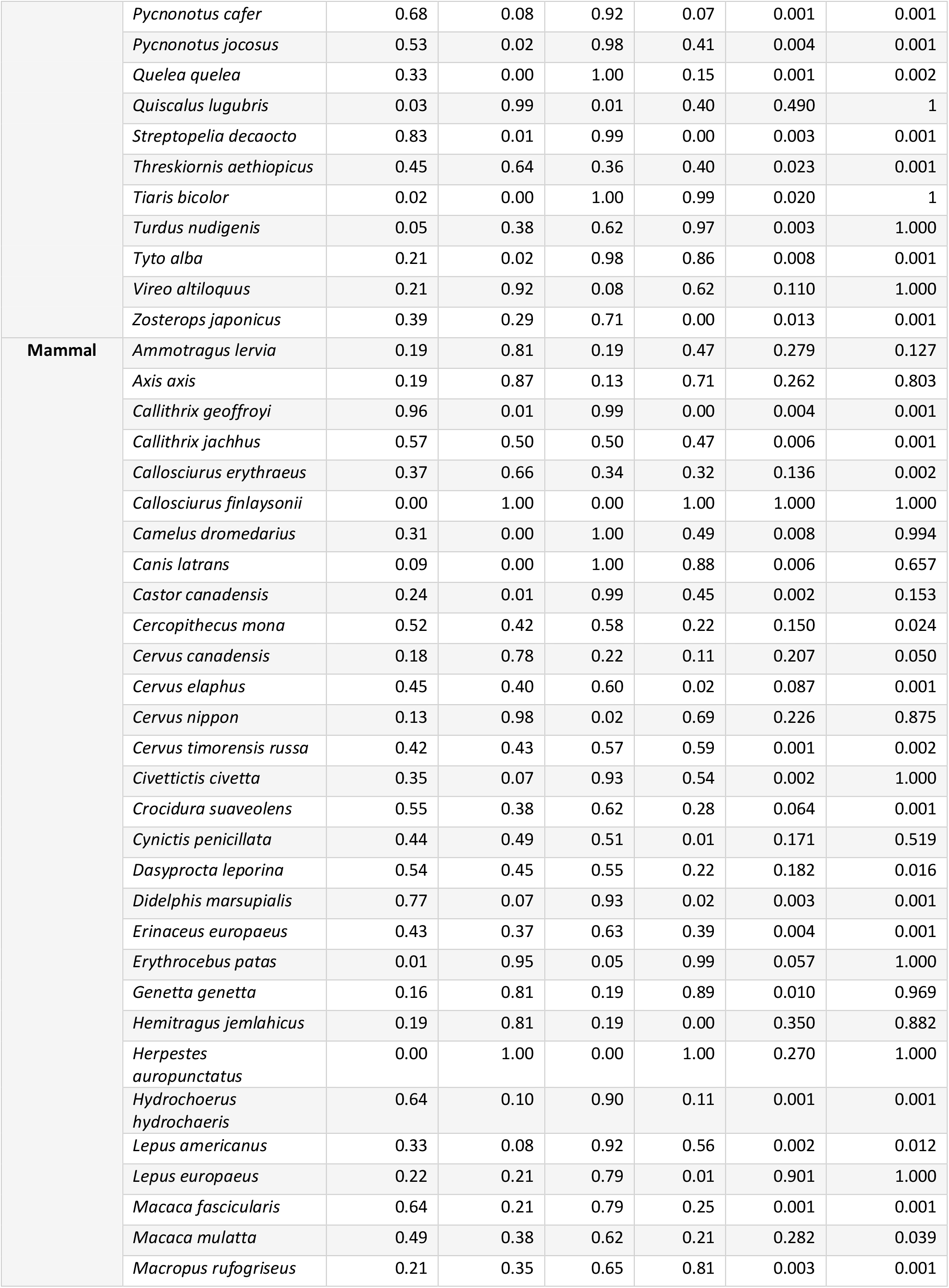
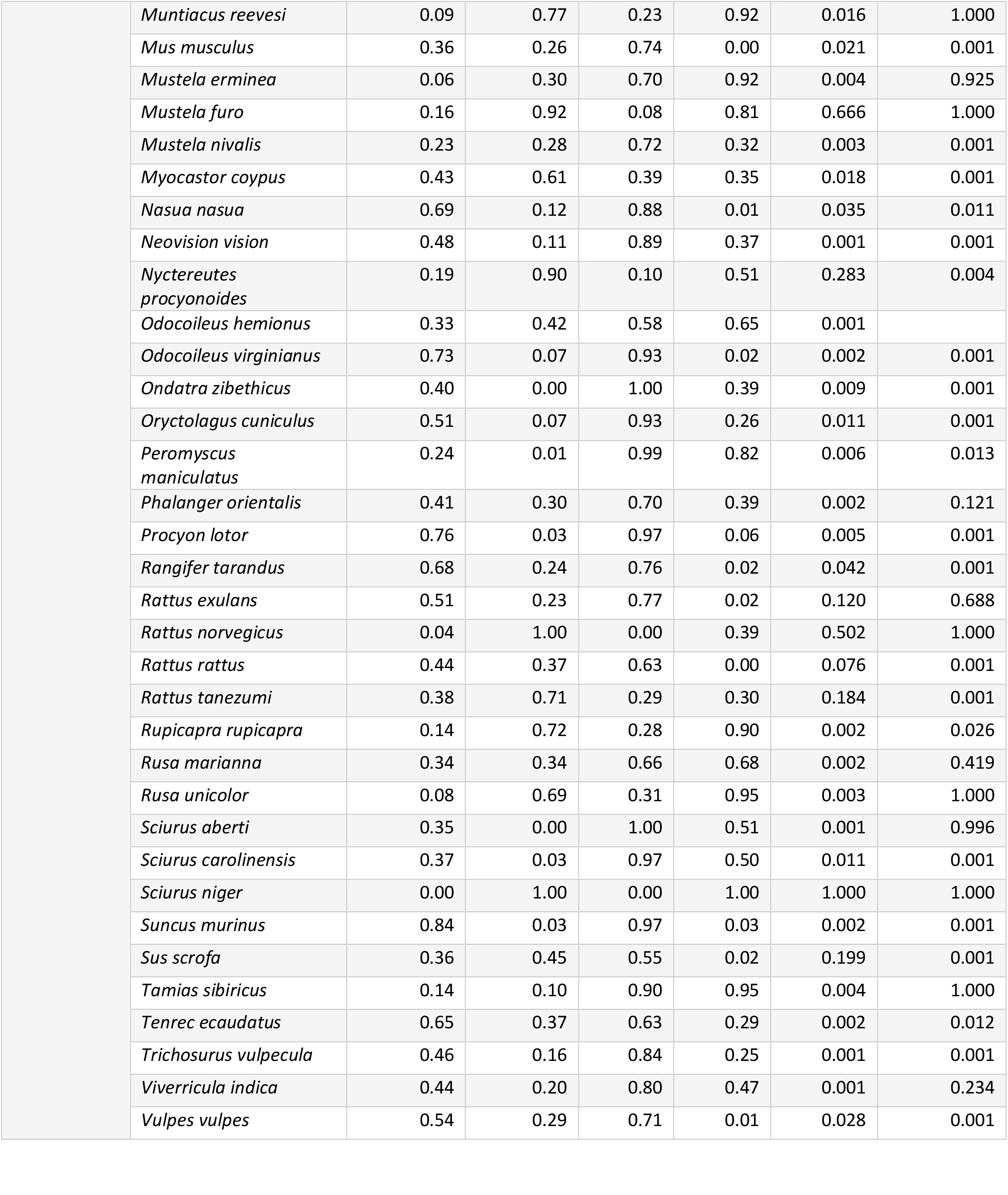
Niche dynamics indices of invasive tetrapods.

| Taxon | Species | Schoener's Index (D) | Expansion (E) | Stability (S) | Unfilling (U) | Similarity | Equivalency |
| --- | --- | --- | --- | --- | --- | --- | --- |
| Amphibian | *Eleutherodactylus coqui* | 0.14 | 0.86 | 0.14 | 0.23 | 0.349 | 1.000 |
|  | *Eleutherodactylus johnstonei* | 0.25 | 0.70 | 0.30 | 0.28 | 0.266 | 1.000 |
|  | *Eleutherodactylus planirostris* | 0.21 | 0.73 | 0.27 | 0.01 | 0.434 | 0.832 |
|  | *Lithobates catesbeianus* | 0.32 | 0.31 | 0.69 | 0.00 | 0.022 | 0.001 |
|  | *Litoria aurea* | 0.25 | 0.82 | 0.18 | 0.59 | 0.146 | 0.444 |
|  | *Osteopilus septentrionalis* | 0.42 | 0.18 | 0.82 | 0.11 | 0.119 | 0.001 |
|  | *Pelophylax bedriagae* | 0.41 | 0.49 | 0.51 | 0.03 | 0.190 | 0.933 |
|  | *Pelophylax kurtumuelleri* | 0.15 | 0.70 | 0.30 | 0.91 | 0.476 | 0.001 |
|  | *Pelophylax perezii* | 0.34 | 0.64 | 0.36 | 0.60 | 0.001 | 0.001 |
|  | *Pelophylax ridibundus* | 0.07 | 0.00 | 1.00 | 0.80 | 0.023 | 1.000 |
|  | *Rhinella marina* | 0.87 | 0.00 | 1.00 | 0.01 | 0.001 | 0.001 |
|  | *Scinax ruber* | 0.09 | 0.00 | 1.00 | 0.81 | 0.003 | 0.300 |
|  | *Scinax x-signatus* | 0.02 | 0.00 | 1.00 | 0.96 | 0.022 | 0.636 |
|  | *Triturus carnifex* | 0.18 | 0.00 | 1.00 | 0.78 | 0.002 | 0.477 |
|  | *Xenopus laevis* | 0.54 | 0.08 | 0.92 | 0.22 | 0.004 | 0.001 |
| Reptile | *Anolis aeneus* | 0.03 | 0.98 | 0.02 | 0.73 | 0.488 | 0.996 |
|  | *Anolis carolinensis* | 0.19 | 0.67 | 0.33 | 0.75 | 0.105 | 0.902 |
|  | *Anolis cristatellus* | 0.24 | 0.53 | 0.47 | 0.82 | 0.002 | 0.924 |
|  | *Anolis equestris* | 0.33 | 0.60 | 0.40 | 0.42 | 0.276 | 0.020 |
|  | *Boiga irregularis* | 0.00 | 0.00 | 1.00 | 1.00 | 0.082 |  |
|  | *Caiman crocodilus* | 0.01 | 0.85 | 0.15 | 1.00 | 0.032 |  |
|  | *Chamaeleo jacksonii* | 0.00 | 1.00 | 0.00 | 1.00 | 0.002 | 0.001 |
|  | *Chamaeleo chamaeleon* | 0.31 | 0.02 | 0.98 | 0.70 | 0.004 | 1.000 |
|  | *Ctenosaura similis* | 0.07 | 0.00 | 1.00 | 0.98 | 0.011 | 0.902 |
|  | *Cyrtopodion scabrum* | 0.41 | 0.45 | 0.55 | 0.18 | 0.170 | 0.906 |
|  | *Elaphe guttata* | 0.18 | 0.92 | 0.08 | 0.93 | 0.012 | 1.000 |
|  | *Eunectes murinus* | 0.13 | 0.16 | 0.84 | 0.91 | 0.001 | 0.250 |
|  | *Eunectes notaeus* | 0.43 | 0.07 | 0.93 | 0.52 | 0.001 | 0.022 |
|  | *Iguana iguana* | 0.35 | 0.02 | 0.98 | 0.28 | 0.006 | 0.001 |
|  | *Natrix maura* | 0.05 | 0.00 | 1.00 | 0.95 | 0.008 | 0.553 |
|  | *Testudo horsfieldii* | 0.11 | 0.92 | 0.08 | 0.93 | 0.259 | 0.997 |
|  | *Trachemys scripta elegans* | 0.24 | 0.68 | 0.32 | 0.04 | 0.401 | 0.998 |
|  | *Tupinambis teguixin* | 0.03 | 0.13 | 0.87 | 0.99 | 0.010 |  |
|  | *Varanus indicus* | 0.05 | 0.00 | 1.00 | 0.84 | 0.006 | 0.802 |
|  | *Varanus niloticus* | 0.06 | 0.10 | 0.90 | 0.97 | 0.009 | 0.167 |
|  | *Zamenis scalaris* | 0.08 | 0.00 | 1.00 | 0.80 | 0.007 | 0.705 |
|  | *Hemidactylus frenatus* | 0.67 | 0.01 | 0.99 | 0.08 | 0.001 | 0.001 |
|  | *Hemidactylus mabouia* | 0.61 | 0.04 | 0.96 | 0.43 | 0.004 | 0.001 |
|  | *Hierophis viridiflavus* | 0.02 | 0.00 | 1.00 | 0.93 | 0.026 |  |
|  | *Podarcis siculus* | 0.24 | 0.78 | 0.22 | 0.87 | 0.002 | 0.207 |
|  | *Python bivittatus* | 0.18 | 0.08 | 0.92 | 0.86 | 0.003 | 1.000 |
|  | *Python regius* | 0.32 | 0.72 | 0.28 | 0.43 | 0.174 | 0.001 |
|  | *Ramphotyphlops*<br>*braminus* | 0.85 | 0.07 | 0.93 | 0.05 | 0.001 | 0.001 |
|  | *Trioceros jacksonii* | 0.16 | 0.85 | 0.15 | 0.94 | 0.002 | 1.000 |
| **Bird** | *Acridotheres fuscus* | 0.61 | 0.22 | 0.78 | 0.09 | 0.002 | 0.001 |
|  | *Acridotheres tristis* | 0.62 | 0.19 | 0.81 | 0.07 | 0.007 | 0.001 |
|  | *Alectoris chukar* | 0.64 | 0.08 | 0.92 | 0.02 | 0.028 | 0.001 |
|  | *Alopochen aegyptiaca* | 0.36 | 0.83 | 0.17 | 0.39 | 0.061 | 0.001 |
|  | *Amazona amazonica* | 0.50 | 0.11 | 0.89 | 0.19 | 0.003 | 0.001 |
|  | *Anser anser* | 0.69 | 0.11 | 0.89 | 0.07 | 0.001 | 0.001 |
|  | *Anser indicus* | 0.18 | 0.20 | 0.80 | 0.90 | 0.020 | 0.001 |
|  | *Branta canadensis* | 0.51 | 0.01 | 0.99 | 0.29 | 0.006 | 0.001 |
|  | *Branta hutchinsii* | 0.25 | 0.02 | 0.98 | 0.85 | 0.015 | 0.001 |
|  | *Carpodacus mexicanus* | 0.47 | 0.11 | 0.89 | 0.55 |  |  |
|  | *Chen canagica* | 0.11 | 0.00 | 1.00 | 0.90 | 0.009 | 1.000 |
|  | *Circus approximans* | 0.38 | 0.04 | 0.96 | 0.44 | 0.006 | 0.001 |
|  | *Columba livia* | 0.80 | 0.01 | 0.99 | 0.01 | 0.003 | 0.001 |
|  | *Corvus splendens* | 0.54 | 0.01 | 0.99 | 0.06 |  |  |
|  | *Cygnus olor* | 0.52 | 0.28 | 0.72 | 0.05 | 0.046 | 0.001 |
|  | *Dicrurus macrocercus* | 0.01 | 0.00 | 1.00 | 1.00 |  |  |
|  | *Estrilda astrild* | 0.43 | 0.44 | 0.56 | 0.02 | 0.074 | 0.001 |
|  | *Gallus gallus* | 0.70 | 0.10 | 0.90 | 0.11 | 0.007 | 0.001 |
|  | *Garrulax canorus* | 0.34 | 0.62 | 0.38 | 0.04 | 0.104 | 0.005 |
|  | *Grus antigone* | 0.34 | 0.78 | 0.22 | 0.57 | 0.459 | 0.112 |
|  | *Gymnorhina tibicen* | 0.22 | 0.03 | 0.97 | 0.74 | 0.008 | 0.001 |
|  | *Horornis diphone* | 0.26 | 0.69 | 0.31 | 0.65 | 0.200 | 0.290 |
|  | *Leiothrix lutea* | 0.38 | 0.51 | 0.49 | 0.43 | 0.001 | 0.001 |
|  | *Molothrus bonariensis* | 0.29 | 0.00 | 1.00 | 0.35 | 0.007 | 0.001 |
|  | *Myiopsitta monachus* | 0.50 | 0.50 | 0.50 | 0.00 | 0.065 | 0.001 |
|  | *Oxyura jamaicensis* | 0.66 | 0.07 | 0.93 | 0.13 | 0.003 | 0.001 |
|  | *Passer domesticus* | 0.77 | 0.02 | 0.98 | 0.01 | 0.012 | 0.001 |
|  | *Pavo cristatus* | 0.60 | 0.30 | 0.70 | 0.07 | 0.010 | 0.001 |
|  | *Phasianus colchicus* | 0.58 | 0.18 | 0.82 | 0.12 | 0.003 | 0.001 |
|  | *Pitangus sulphuratus* | 0.04 | 0.00 | 1.00 | 0.87 | 0.015 | 0.999 |
|  | *Ploceus cucullatus* | 0.79 | 0.06 | 0.94 | 0.16 | 0.001 | 0.001 |
|  | *Porphyrio porphyrio* | 0.38 | 0.04 | 0.96 | 0.40 | 0.005 | 0.001 |
|  | *Psittacula krameri* | 0.35 | 0.55 | 0.45 | 0.38 | 0.006 | 0.001 |
|  | *Pycnonotus cafer* | 0.68 | 0.08 | 0.92 | 0.07 | 0.001 | 0.001 |
|  | *Pycnonotus jocosus* | 0.53 | 0.02 | 0.98 | 0.41 | 0.004 | 0.001 |
|  | *Quelea quelea* | 0.33 | 0.00 | 1.00 | 0.15 | 0.001 | 0.002 |
|  | *Quiscalus lugubris* | 0.03 | 0.99 | 0.01 | 0.40 | 0.490 | 1 |
|  | *Streptopelia decaocto* | 0.83 | 0.01 | 0.99 | 0.00 | 0.003 | 0.001 |
|  | *Threskiornis aethiopicus* | 0.45 | 0.64 | 0.36 | 0.40 | 0.023 | 0.001 |
|  | *Tiaris bicolor* | 0.02 | 0.00 | 1.00 | 0.99 | 0.020 | 1 |
|  | *Turdus nudigenis* | 0.05 | 0.38 | 0.62 | 0.97 | 0.003 | 1.000 |
|  | *Tyto alba* | 0.21 | 0.02 | 0.98 | 0.86 | 0.008 | 0.001 |
|  | *Vireo altiloquus* | 0.21 | 0.92 | 0.08 | 0.62 | 0.110 | 1.000 |
|  | *Zosterops japonicus* | 0.39 | 0.29 | 0.71 | 0.00 | 0.013 | 0.001 |
| **Mammal** | *Ammotragus lervia* | 0.19 | 0.81 | 0.19 | 0.47 | 0.279 | 0.127 |
|  | *Axis axis* | 0.19 | 0.87 | 0.13 | 0.71 | 0.262 | 0.803 |
|  | *Callithrix geoffroyi* | 0.96 | 0.01 | 0.99 | 0.00 | 0.004 | 0.001 |
|  | *Callithrix jachhus* | 0.57 | 0.50 | 0.50 | 0.47 | 0.006 | 0.001 |
|  | *Callosciurus erythraeus* | 0.37 | 0.66 | 0.34 | 0.32 | 0.136 | 0.002 |
|  | *Callosciurus finlaysonii* | 0.00 | 1.00 | 0.00 | 1.00 | 1.000 | 1.000 |
|  | *Camelus dromedarius* | 0.31 | 0.00 | 1.00 | 0.49 | 0.008 | 0.994 |
|  | *Canis latrans* | 0.09 | 0.00 | 1.00 | 0.88 | 0.006 | 0.657 |
|  | *Castor canadensis* | 0.24 | 0.01 | 0.99 | 0.45 | 0.002 | 0.153 |
|  | *Cercopithecus mona* | 0.52 | 0.42 | 0.58 | 0.22 | 0.150 | 0.024 |
|  | *Cervus canadensis* | 0.18 | 0.78 | 0.22 | 0.11 | 0.207 | 0.050 |
|  | *Cervus elaphus* | 0.45 | 0.40 | 0.60 | 0.02 | 0.087 | 0.001 |
|  | *Cervus nippon* | 0.13 | 0.98 | 0.02 | 0.69 | 0.226 | 0.875 |
|  | *Cervus timorensis russa* | 0.42 | 0.43 | 0.57 | 0.59 | 0.001 | 0.002 |
|  | *Civettictis civetta* | 0.35 | 0.07 | 0.93 | 0.54 | 0.002 | 1.000 |
|  | *Crocidura suaveolens* | 0.55 | 0.38 | 0.62 | 0.28 | 0.064 | 0.001 |
|  | *Cynictis penicillata* | 0.44 | 0.49 | 0.51 | 0.01 | 0.171 | 0.519 |
|  | *Dasyprocta leporina* | 0.54 | 0.45 | 0.55 | 0.22 | 0.182 | 0.016 |
|  | *Didelphis marsupialis* | 0.77 | 0.07 | 0.93 | 0.02 | 0.003 | 0.001 |
|  | *Erinaceus europaeus* | 0.43 | 0.37 | 0.63 | 0.39 | 0.004 | 0.001 |
|  | *Erythrocebus patas* | 0.01 | 0.95 | 0.05 | 0.99 | 0.057 | 1.000 |
|  | *Genetta genetta* | 0.16 | 0.81 | 0.19 | 0.89 | 0.010 | 0.969 |
|  | *Hemitragus jemlahicus* | 0.19 | 0.81 | 0.19 | 0.00 | 0.350 | 0.882 |
|  | *Herpestes auropunctatus* | 0.00 | 1.00 | 0.00 | 1.00 | 0.270 | 1.000 |
|  | *Hydrochoerus hydrochaeris* | 0.64 | 0.10 | 0.90 | 0.11 | 0.001 | 0.001 |
|  | *Lepus americanus* | 0.33 | 0.08 | 0.92 | 0.56 | 0.002 | 0.012 |
|  | *Lepus europaeus* | 0.22 | 0.21 | 0.79 | 0.01 | 0.901 | 1.000 |
|  | *Macaca fascicularis* | 0.64 | 0.21 | 0.79 | 0.25 | 0.001 | 0.001 |
|  | *Macaca mulatta* | 0.49 | 0.38 | 0.62 | 0.21 | 0.282 | 0.039 |
|  | *Macropus rufogriseus* | 0.21 | 0.35 | 0.65 | 0.81 | 0.003 | 0.001 |

|  |  |  |  |  |  |  |
| --- | --- | --- | --- | --- | --- | --- |
| *Muntiacus reevesi* | 0.09 | 0.77 | 0.23 | 0.92 | 0.016 | 1.000 |
| *Mus musculus* | 0.36 | 0.26 | 0.74 | 0.00 | 0.021 | 0.001 |
| *Mustela erminea* | 0.06 | 0.30 | 0.70 | 0.92 | 0.004 | 0.925 |
| *Mustela furo* | 0.16 | 0.92 | 0.08 | 0.81 | 0.666 | 1.000 |
| *Mustela nivalis* | 0.23 | 0.28 | 0.72 | 0.32 | 0.003 | 0.001 |
| *Myocastor coypus* | 0.43 | 0.61 | 0.39 | 0.35 | 0.018 | 0.001 |
| *Nasua nasua* | 0.69 | 0.12 | 0.88 | 0.01 | 0.035 | 0.011 |
| *Neovison vision* | 0.48 | 0.11 | 0.89 | 0.37 | 0.001 | 0.001 |
| *Nyctereutes*<br>*procyonoides* | 0.19 | 0.90 | 0.10 | 0.51 | 0.283 | 0.004 |
| *Odocoileus hemionus* | 0.33 | 0.42 | 0.58 | 0.65 | 0.001 |  |
| *Odocoileus virginianus* | 0.73 | 0.07 | 0.93 | 0.02 | 0.002 | 0.001 |
| *Ondatra zibethicus* | 0.40 | 0.00 | 1.00 | 0.39 | 0.009 | 0.001 |
| *Oryctolagus cuniculus* | 0.51 | 0.07 | 0.93 | 0.26 | 0.011 | 0.001 |
| *Peromyscus*<br>*maniculatus* | 0.24 | 0.01 | 0.99 | 0.82 | 0.006 | 0.013 |
| *Phalanger orientalis* | 0.41 | 0.30 | 0.70 | 0.39 | 0.002 | 0.121 |
| *Procyon lotor* | 0.76 | 0.03 | 0.97 | 0.06 | 0.005 | 0.001 |
| *Rangifer tarandus* | 0.68 | 0.24 | 0.76 | 0.02 | 0.042 | 0.001 |
| *Rattus exulans* | 0.51 | 0.23 | 0.77 | 0.02 | 0.120 | 0.688 |
| *Rattus norvegicus* | 0.04 | 1.00 | 0.00 | 0.39 | 0.502 | 1.000 |
| *Rattus rattus* | 0.44 | 0.37 | 0.63 | 0.00 | 0.076 | 0.001 |
| *Rattus tanezumi* | 0.38 | 0.71 | 0.29 | 0.30 | 0.184 | 0.001 |
| *Rupicapra rupicapra* | 0.14 | 0.72 | 0.28 | 0.90 | 0.002 | 0.026 |
| *Rusa marianna* | 0.34 | 0.34 | 0.66 | 0.68 | 0.002 | 0.419 |
| *Rusa unicolor* | 0.08 | 0.69 | 0.31 | 0.95 | 0.003 | 1.000 |
| *Sciurus aberti* | 0.35 | 0.00 | 1.00 | 0.51 | 0.001 | 0.996 |
| *Sciurus carolinensis* | 0.37 | 0.03 | 0.97 | 0.50 | 0.011 | 0.001 |
| *Sciurus niger* | 0.00 | 1.00 | 0.00 | 1.00 | 1.000 | 1.000 |
| *Suncus murinus* | 0.84 | 0.03 | 0.97 | 0.03 | 0.002 | 0.001 |
| *Sus scrofa* | 0.36 | 0.45 | 0.55 | 0.02 | 0.199 | 0.001 |
| *Tamias sibiricus* | 0.14 | 0.10 | 0.90 | 0.95 | 0.004 | 1.000 |
| *Tenrec ecaudatus* | 0.65 | 0.37 | 0.63 | 0.29 | 0.002 | 0.012 |
| *Trichosurus vulpecula* | 0.46 | 0.16 | 0.84 | 0.25 | 0.001 | 0.001 |
| *Viverricula indica* | 0.44 | 0.20 | 0.80 | 0.47 | 0.001 | 0.234 |
| *Vulpes vulpes* | 0.54 | 0.29 | 0.71 | 0.01 | 0.028 | 0.001 |

**Table 3:**
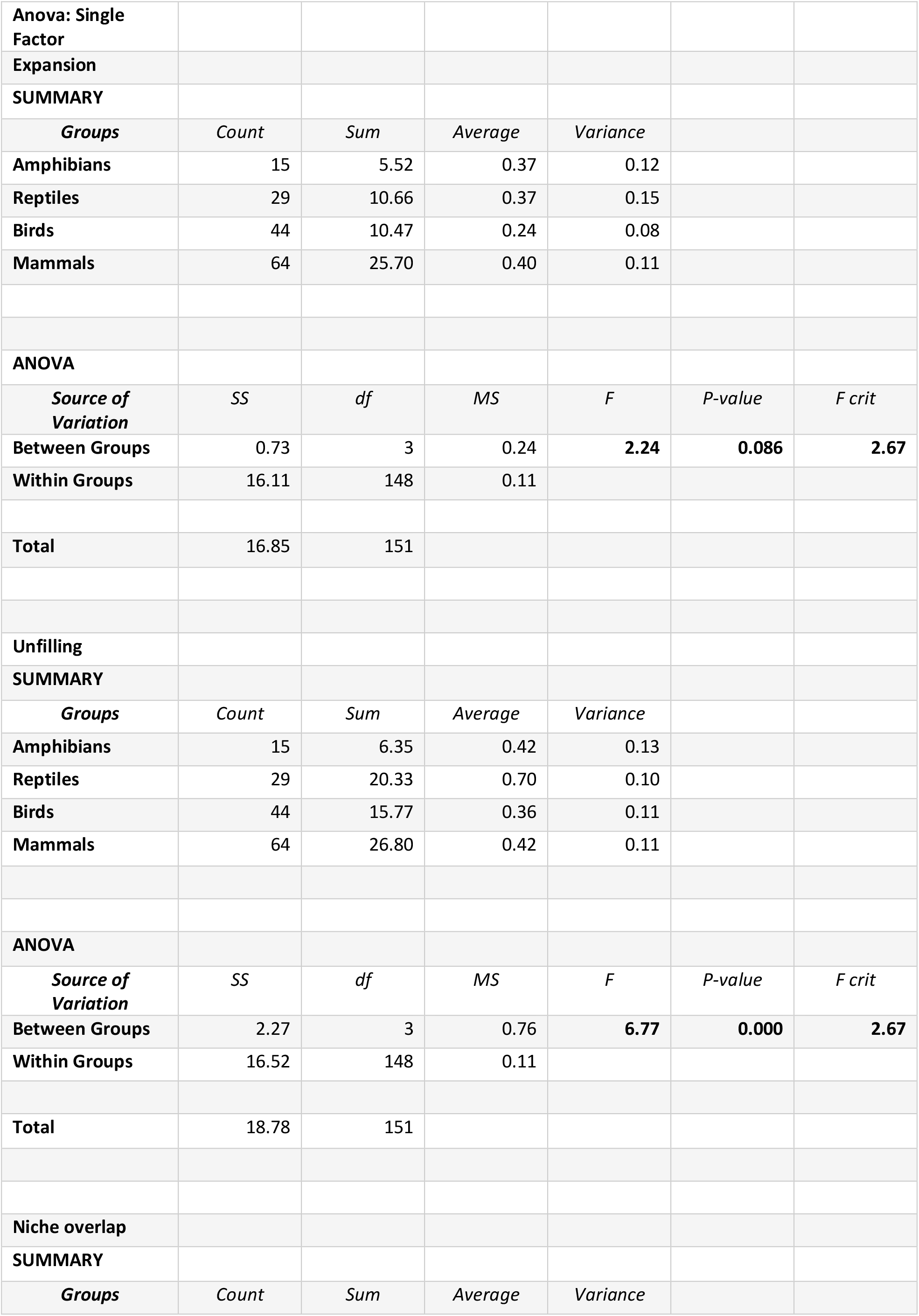
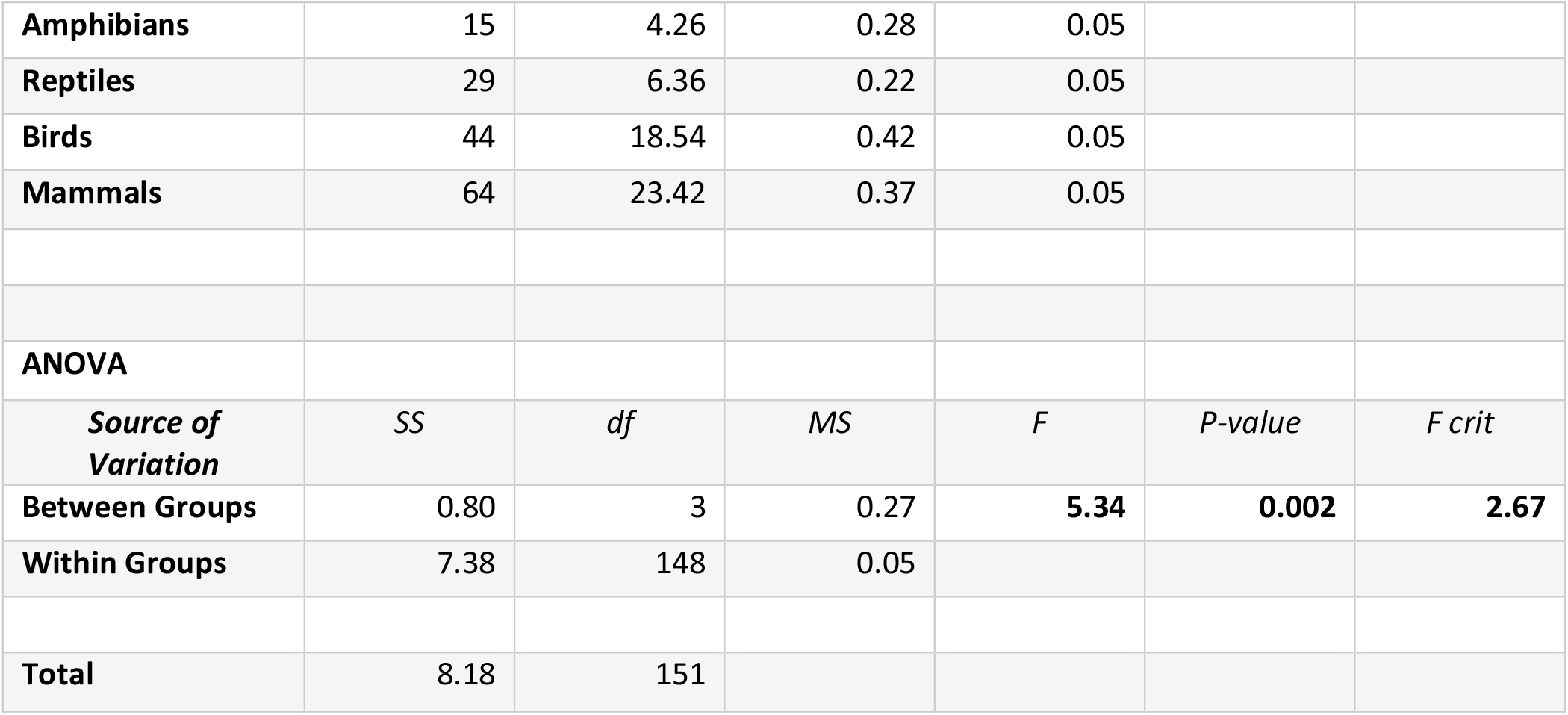
Analysis of Variance of niche expansion, unfilling, and overlap across tetrapod.

|  |  |  |  |  |  |  |
| --- | --- | --- | --- | --- | --- | --- |
| **Anova: Single Factor** |  |  |  |  |  |  |
| **Expansion** |  |  |  |  |  |  |
| **SUMMARY** |  |  |  |  |  |  |
| *Groups* | *Count* | *Sum* | *Average* | *Variance* |  |  |
| **Amphibians** | 15 | 5.52 | 0.37 | 0.12 |  |  |
| **Reptiles** | 29 | 10.66 | 0.37 | 0.15 |  |  |
| **Birds** | 44 | 10.47 | 0.24 | 0.08 |  |  |
| **Mammals** | 64 | 25.70 | 0.40 | 0.11 |  |  |
| **ANOVA** |  |  |  |  |  |  |
| *Source of Variation* | *SS* | *df* | *MS* | *F* | *P-value* | *F crit* |
| **Between Groups** | 0.73 | 3 | 0.24 | **2.24** | **0.086** | **2.67** |
| **Within Groups** | 16.11 | 148 | 0.11 |  |  |  |
| **Total** | 16.85 | 151 |  |  |  |  |
| **Unfilling** |  |  |  |  |  |  |
| **SUMMARY** |  |  |  |  |  |  |
| *Groups* | *Count* | *Sum* | *Average* | *Variance* |  |  |
| **Amphibians** | 15 | 6.35 | 0.42 | 0.13 |  |  |
| **Reptiles** | 29 | 20.33 | 0.70 | 0.10 |  |  |
| **Birds** | 44 | 15.77 | 0.36 | 0.11 |  |  |
| **Mammals** | 64 | 26.80 | 0.42 | 0.11 |  |  |
| **ANOVA** |  |  |  |  |  |  |
| *Source of Variation* | *SS* | *df* | *MS* | *F* | *P-value* | *F crit* |
| **Between Groups** | 2.27 | 3 | 0.76 | **6.77** | **0.000** | **2.67** |
| **Within Groups** | 16.52 | 148 | 0.11 |  |  |  |
| **Total** | 18.78 | 151 |  |  |  |  |
| **Niche overlap** |  |  |  |  |  |  |
| **SUMMARY** |  |  |  |  |  |  |
| *Groups* | *Count* | *Sum* | *Average* | *Variance* |  |  |
| **Amphibians** | 15 | 4.26 | 0.28 | 0.05 |  |  |
| **Reptiles** | 29 | 6.36 | 0.22 | 0.05 |  |  |
| **Birds** | 44 | 18.54 | 0.42 | 0.05 |  |  |
| **Mammals** | 64 | 23.42 | 0.37 | 0.05 |  |  |
| **ANOVA** |  |  |  |  |  |  |
| ***Source of Variation*** | ***SS*** | ***df*** | ***MS*** | ***F*** | ***P-value*** | ***F crit*** |
| **Between Groups** | 0.80 | 3 | 0.27 | **5.34** | **0.002** | **2.67** |
| **Within Groups** | 7.38 | 148 | 0.05 |  |  |  |
| **Total** | 8.18 | 151 |  |  |  |  |

**Figure 2:**
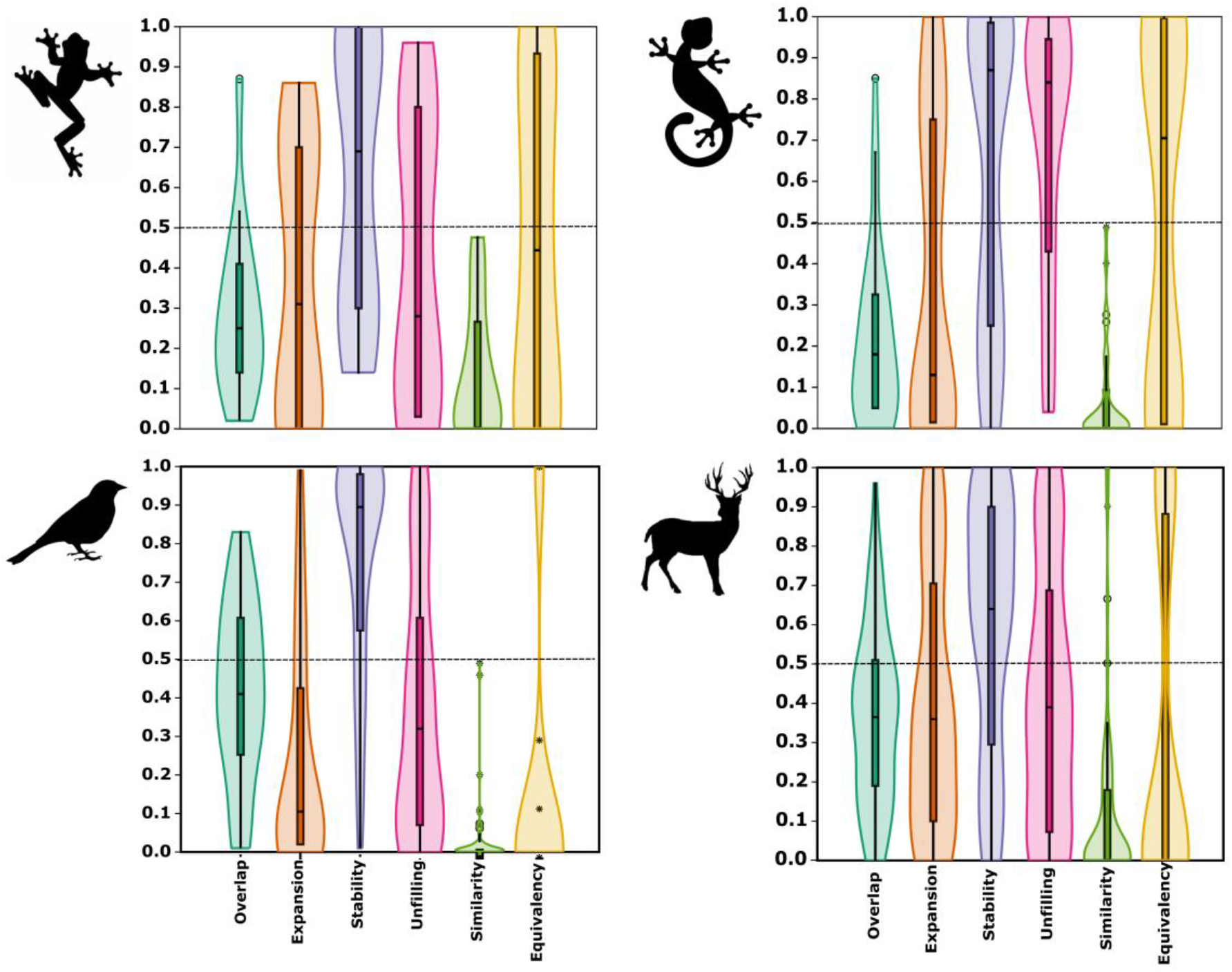
Niche dynamics indices across tetrapods: niche overlap, expansion, stability, unfilling, similarity test and equivalency test. The violin plot shows the distribution of niche dynamics indices across all tetrapod taxa.

**Figure 3:**
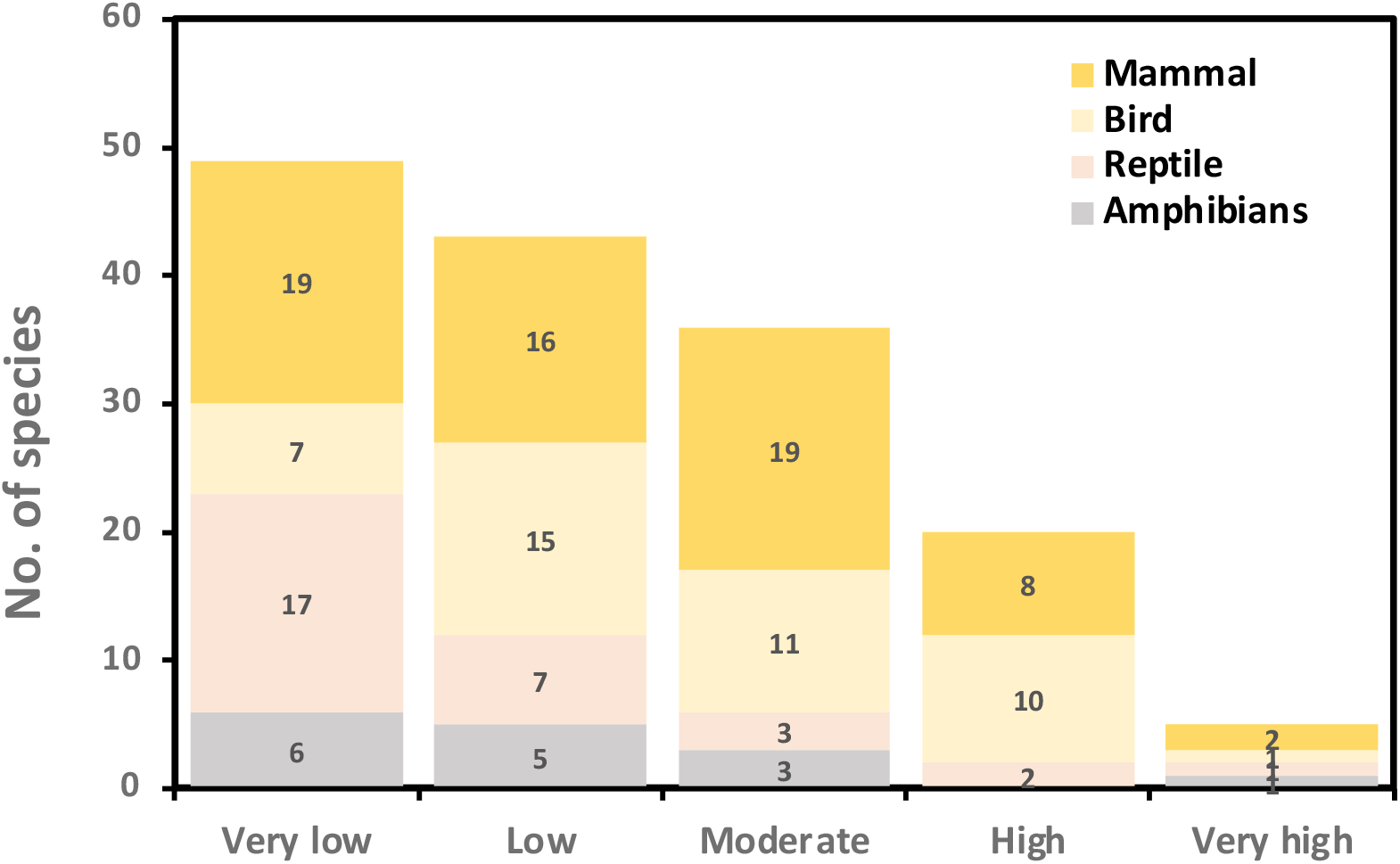
Niche overlap index (Schoener’s D) across tetrapod taxa. The value ranges from zero to 1, and it is categorised as very low (zero to 0.2), low (0.2 to 0.4), moderate (0.4 to 0.6), high (0.6 to 0.8), and very high (0.8 to 1).

## Discussion

This study presents the global-level assessment of the niche dynamics of 152 invasive tetrapods. There is inherent complexity of invasion processes as the invasion dynamics are influenced by diverse biotic and abiotic factors, including species interactions, native biodiversity, ecosystem services, and human-mediated factors like migration (Hulme, 2009; Gallardo and Aldridge, 2013), Due to the impracticality of comprehensively understanding biotic interactions for over 150 species across various taxa, our focus was primarily on utilising bioclimatic variables, considering the generalist nature of invasive species, to investigate whether tetrapods exhibit niche conservatism or show niche shift. The phenomenon of both niche shifts and conservatism has been documented in the literature. For instance, some introduced birds and mammals in Europe have demonstrated niche conservatism (Strubbe et al., 2013; Sales et al., 2016). The study by Strubbe et al. (2015) highlighted certain non-native vertebrates that maintain their niches in the introduced region. One could argue that niche conservatism is a primary mechanism through which a species is able to expand into various regions worldwide (Peterson, 2011; Pyron et al., 2015; Strubbe et al., 2015). Expanding on this body of knowledge, our study extends the niche conservatism test to a larger cohort of invasive vertebrates, revealing a prevalence of niche shifts in most of the tetrapods except for invasive birds. This pattern aligns with similar findings in invasive mammals (Mahapatra et al., under review). Conversely, invasive terrestrial plants exhibit niche conservatism (Petitpierre et al., 2012), while invasive reptiles adhere to niche conservatism (Aravind et al., 2023); this was shown in the present study as well. The nuanced dynamics of niche shifts are evident in the case of introduced plants (Atwater et al., 2018) and the majority of invasive freshwater molluscs (Mahapatra et al., 2023), which exhibit niche shifts in their introduced ranges. Contrasting results have been reported, with Liu et al. (2020) indicating that many invasive species adhere to niche conservatism, while Tingley et al. (2014) found evidence of niche expansion in the cane toad (*Rhinella marina*) in Australia. In the case of non-native vertebrates, like wild boar, niche shifts have been observed (Strubbe et al., 2013, 2015). The niche shift can be explained by the unfilling, which means the presence of an introduced environment analogous to the native region (Sales et al. 2016).

The variability in outcomes across studies can be attributed to different metrics used to measure niche shifts and the absence of clear thresholds for determining niche conservatism versus niche shift (Bates and Bertelsmeier, 2021). Interpretation biases and subjective beliefs further contribute to disparate conclusions about niche shifts (Bates and Bertelsmeier, 2021). While most studies reject the niche conservatism hypothesis for invasive species, our global review of tetrapods demonstrates a high degree of niche expansion, particularly in non-bird taxa. Birds, however, exhibit niches in the introduced ranges that closely mirror their native niches or home away from home. Notably, aquatic species and those introduced more recently exhibit lower niche similarity, a finding consistent with Liu et al. (2020). Despite the global review suggesting limited niche expansion between native and introduced ranges, our study on tetrapods reveals substantial niche expansion, with amphibians, often breeding in water, showing a significant number of species with low (<0.4) niche overlap between native and invaded ranges, indicative of niche shift. Aravind et al. (2022) showed that 90% of invasive species move to similar habitats, akin to “home away from home,” while only 10% exhibit niche shifts or “finding a new home,” a trend also echoed in other studies (Liu et al., 2020). The existing body of literature is biased towards studies in the global north, focusing primarily on single species between two regions rather than offering a comprehensive global-level assessment (Liu et al., 2020).

In conclusion, our global assessment of the niche dynamics of 152 invasive tetrapods reveals a nuanced interplay between niche shifts and conservatism. While birds often retain their native niches in introduced ranges, non-bird taxa, particularly amphibians, exhibit substantial niche expansion. The variability in outcomes across taxa underscores the complexity of invasion dynamics and the need for nuanced approaches to understanding species responses. The contrasting patterns observed highlight the importance of taxon-specific analyses. Our findings contribute to the ongoing discourse on the role of niche dynamics in biological invasions, emphasising the necessity for a comprehensive and global perspective to enhance our understanding of invasive species’ ecological adaptability.

## References

Aravind, N.A., Bhat, H.N. P. and Mahapatra, B.B. Niche shifts, low haplotype diversity and invasion potentials of invasive snail Lissachatina fulica (Gastropoda: Achatinidae), Biological Invasions (Accepted)

Aravind N.A., M. Uma Shaanker, H.N.P. Bhat, B. Charles, R. Uma Shaanker, M.A. Shah and G. Ravikanth. 2022. Niche shift in Invasive species: Is it simply a case of “home away from home” or finding a “new home”? Biodiversity and Conservation 31: 2625–2638

Aravind, N.A., Mohopatra, P.P., Bhat, H.P. and Narayanan, S., 2023. Pets or predators? climate change and invasion risk of red-eared slider (Trachemys scripta elgans). Records of the Zoological Survey of India, pp.185–197.

Atwater, D.Z., Ervine, C. and Barney, J.N., 2018. Climatic niche shifts are common in introduced plants. Nature Ecology & Evolution, 2(1), pp.34–43.

Bates, O.K., Ollier, S. and Bertelsmeier, C., 2020. Smaller climatic niche shifts in invasive than non-invasive alien ant species. Nature Communications, 11(1), p.5213.

Bates, O.K. and Bertelsmeier, C., 2021. Climatic niche shifts in introduced species. Current Biology, 31(19), pp.R1252–R1266.

Bellard, C., Thuiller, W., Leroy, B., Genovesi, P., Bakkenes, M. and Courchamp, F., 2013. Will climate change promote future invasions?. Global Change Biology, 19(12), pp.3740–3748.

Broennimann, O., Treier, U.A., Müller‐Schärer, H., Thuiller, W., Peterson, A.T. and Guisan, A., 2007. Evidence of climatic niche shift during biological invasion. Ecology Letters, 10(8), pp.701–709.

Broennimann, O., Fitzpatrick, M.C., Pearman, P.B., Petitpierre, B., Pellissier, L., Yoccoz, N.G., Thuiller, W., Fortin, M.J., Randin, C., Zimmermann, N.E. and Graham, C.H., 2012. Measuring ecological niche overlap from occurrence and spatial environmental data. Global Ecology and Biogeography, 21(4), pp.481–497.

Broennimann, O., Petitpierre, B., Chevalier, M., González-Suárez, M., Jeschke, J.M., Rolland, J., Gray, S.M., Bacher, S. and Guisan, A., 2021. Distance to native climatic niche margins explains establishment success of alien mammals. Nature Communications, 12(1), p.2353.

Brondizio, E., Díaz, S.M., Settele, J., Ngo, H., Gueze, M., Aumeeruddy-Thomas, Y., Bai, X., Geschke, A., Molnár, Z., Niamir, A. and Pascual, U., 2019. Assessing a planet in transformation: Rationale and approach of the IPBES Global Assessment on Biodiversity and Ecosystem Services.

Diez, J.M., Hulme, P.E. and Duncan, R.P., 2012. Using prior information to build probabilistic invasive species risk assessments. Biological Invasions, 14, pp.681–691.

Early, R., Bradley, B.A., Dukes, J.S., Lawler, J.J., Olden, J.D., Blumenthal, D.M., Gonzalez, P., Grosholz, E.D., Ibañez, I., Miller, L.P. and Sorte, C.J., 2016. Global threats from invasive alien species in the twenty-first century and national response capacities. Nature Communications, 7(1), p.12485.

Early, R. and Sax, D.F., 2014. Climatic niche shifts between species’ native and naturalized ranges raise concern for ecological forecasts during invasions and climate change. Global Ecology and Biogeography, 23(12), pp.1356–1365.

Fick, S.E. and Hijmans, R.J., 2017. WorldClim 2: new 1‐km spatial resolution climate surfaces for global land areas. International Journal of Climatology, 37(12), pp.4302–4315.

Gallardo, B. and Aldridge, D.C., 2013. The ‘dirty dozen’: socio‐economic factors amplify the invasion potential of 12 high‐risk aquatic invasive species in Great Britain and Ireland. Journal of Applied Ecology, 50(3), pp.757–766.

Gallardo, B., Bacher, S., Bradley, B., Comín, F.A., Gallien, L., Jeschke, J.M., Sorte, C.J. and Vilà, M., 2019. InvasiBES: Understanding and managing the impacts of Invasive alien species on Biodiversity and Ecosystem Services. NeoBiota, 50, pp.109–122.

Guisan, A., Petitpierre, B., Broennimann, O., Daehler, C. and Kueffer, C., 2014. Unifying niche shift studies: insights from biological invasions. Trends in Ecology & Evolution, 29(5), pp.260–269.

Hulme, P.E., 2009. Trade, transport and trouble: managing invasive species pathways in an era of globalization. Journal of Applied Ecology, 46(1), pp.10–18.

Liu, C., Wolter, C., Xian, W. and Jeschke, J.M., 2020. Most invasive species largely conserve their climatic niche. Proceedings of the National Academy of Sciences, 117(38), pp.23643–23651.

Lockwood, J.L., Hoopes, M.F. and Marchetti, M.P., 2013. Invasion Ecology. John Wiley & Sons.

Mahapatra, B.B., Das, N.K., Jadhav, A., Roy, A. and Aravind, N.A., 2023. Global freshwater mollusc invasion: pathways, potential distribution, and niche shift. Hydrobiologia, pp.1–20.

Mahapatra, B.B. G, Ravikanth Aravind N.A., Potential distribution and niche dynamics of invasive mammals. Ecological Complexity (Manuscript under review)

Manel, S., Williams, H.C. and Ormerod, S.J., 2001. Evaluating presence–absence models in ecology: the need to account for prevalence. Journal of Applied Ecology, 38(5), pp.921–931.

Peterson, A.T., 2011. Ecological niche conservatism: A time‐structured review of evidence. Journal of Biogeography, 38(5), pp.817–827.

Peterson, A.T., Stewart, A., Mohamed, K.I. and Araújo, M.B., 2008. Shifting global invasive potential of European plants with climate change. PLoS One, 3(6), p.e2441.

Petitpierre, B., Kueffer, C., Broennimann, O., Randin, C., Daehler, C. and Guisan, A., 2012. Climatic niche shifts are rare among terrestrial plant invaders. Science, 335(6074), pp.1344–1348.

Pyron, R.A., Costa, G.C., Patten, M.A. and Burbrink, F.T., 2015. Phylogenetic niche conservatism and the evolutionary basis of ecological speciation. Biological Reviews, 90(4), pp.1248–1262.

Pyšek, P., Hulme, P.E., Simberloff, D., Bacher, S., Blackburn, T.M., Carlton, J.T., Dawson, W., Essl, F., Foxcroft, L.C., Genovesi, P. and Jeschke, J.M., 2020. Scientists’ warning on invasive alien species. Biological Reviews, 95(6), pp.1511–1534.

Ribeiro, J., Reino, L., Schindler, S., Strubbe, D., Vall-llosera, M., Araújo, M.B., Capinha, C., Carrete, M., Mazzoni, S., Monteiro, M. and Moreira, F., 2019. Trends in legal and illegal trade of wild birds: A global assessment based on expert knowledge. Biodiversity and Conservation, 28, pp.3343–3369.

Rödder, D. and Engler, J.O., 2011. Quantitative metrics of overlaps in Grinnellian niches: advances and possible drawbacks. Global Ecology and Biogeography, 20(6): 915–927.

Sales LP, Ribeiro BR, Hayward MW, Paglia A, Passamani M, Loyola R. Niche conservatism and the invasive potential of the wild boar. Journal of Animal Ecology. 2017 Sep;86(5):1214–1223. doi: 10.1111/1365-2656.12721.

Seebens, H., Blackburn, T.M., Dyer, E.E., Genovesi, P., Hulme, P.E., Jeschke, J.M., Pagad, S., Pyšek, P., Winter, M., Arianoutsou, M. and Bacher, S., 2017. No saturation in the accumulation of alien species worldwide. Nature communications, 8(1), p.14435.

Seebens, H., Blackburn, T.M., Hulme, P.E., Van Kleunen, M., Liebhold, A.M., Orlova‐Bienkowskaja, M., Pyšek, P., Schindler, S. and Essl, F., 2021. Around the world in 500 years: Inter‐regional spread of alien species over recent centuries. Global Ecology and Biogeography, 30(8), pp.1621–1632.

Stroud, J.T., 2021. Island species experience higher niche expansion and lower niche conservatism during invasion. Proceedings of the National Academy of Sciences, 118(1), p.e2018949118.

Strubbe, D., Beauchard, O. and Matthysen, E., 2015. Niche conservatism among non‐native vertebrates in Europe and North America. Ecography, 38(3): 321–329.

Strubbe, D., Broennimann, O., Chiron, F. and Matthysen, E., 2013. Niche conservatism in non‐native birds in Europe: niche unfilling rather than niche expansion. Global Ecology and Biogeography, 22(8): 962–970.

Strubbe, D. and Matthysen, E., 2009. Predicting the potential distribution of invasive ring-necked parakeets Psittacula krameri in northern Belgium using an ecological niche modelling approach. Biological Invasions, 11: 497–513.

Swets, J.A., 1988. Measuring the accuracy of diagnostic systems. Science, 240(4857): 1285–1293.

Team, R.C., 2018. A language and environment for statistical computing. R Foundation for Statistical Computing, Vienna, Austria. Available online: https://www.R-project.org/ (accessed on 11 September 2020).

Tingley, R., Vallinoto, M., Sequeira, F. and Kearney, M.R., 2014. Realized niche shift during a global biological invasion. Proceedings of the National Academy of Sciences, 111(28): 10233–10238.

Wiens, J.J. and Graham, C.H., 2005. Niche conservatism: integrating evolution, ecology, and conservation biology. Annual Review of. Ecology, Evolution and. Systematics, 36: 519–539.

Yang, R., Cao, R., Gong, X. and Feng, J., 2023. Large shifts of niche and range in the golden apple snail (Pomacea canaliculata), an aquatic invasive species. Ecosphere, 14(1), p.e4391.

